# Selective attention to audiovisual speech routes activity through recurrent feedback-feedforward loops between different nodes of the speech network

**DOI:** 10.1101/2023.07.17.549287

**Authors:** Patrik Wikman, Viljami Salmela, Eetu Sjöblom, Miika Leminen, Matti Laine, Kimmo Alho

## Abstract

Selective attention related top-down modulation plays a significant role in separating relevant speech from irrelevant background speech when vocal attributes separating concurrent speakers are small and continuously evolving. Electrophysiological studies have shown that such top-down modulation enhances neural tracking of attended speech. Yet, the specific cortical regions involved remain unclear due to the limited spatial resolution of most electrophysiological techniques. To overcome such limitations, we collected both EEG (high temporal resolution) and fMRI (high spatial resolution), while human participants selectively attended to speakers in audiovisual scenes containing overlapping cocktail party speech. To utilize the advantages of the respective techniques, we analysed neural tracking of speech using the EEG data and performed representational dissimilarity-based EEG-fMRI fusion. We observed that attention enhanced neural tracking and modulated EEG correlates throughout the latencies studied. Further, attention related enhancement of neural tracking fluctuated in predictable temporal profiles. We discuss how such temporal dynamics could arise from a combination of interactions between attention and prediction as well as plastic properties of the auditory cortex. EEG-fMRI fusion revealed attention related iterative feedforward-feedback loops between hierarchically organised nodes of the ventral auditory object related processing stream. Our findings support models where attention facilitates dynamic neural changes in the auditory cortex, ultimately aiding discrimination of relevant sounds from irrelevant ones while conserving neural resources.

## Introduction

Humans effortlessly recognize and separate auditory objects in complex sound environments. This ability relies on hierarchical neural processing in the auditory ventral “what” stream, where sequential processing stages extract and integrate increasingly complex object attributes[1, 2] – starting with processing of simple features (e.g. frequency) in the primary auditory cortex, progressing to complex acoustic structures (e.g., frequency-modulated sweeps) in secondary areas and selectivity for complete auditory objects in the anterior superior temporal cortex. [3-6] The ventral stream terminates in the anterior temporal and inferior frontal cortex where sound category and semantic information is apparently stored. [7-9].

In the absence of spatial cues, there are usually only subtle differences in the vocal attributes that separate concurrent speakers from each other. [10] Therefore, top-down modulation facilitated by selective attention plays a significant role in separating relevant speech objects from irrelevant background speech. [10, 11] This top-down modulation is classically assumed to enhance the gain [12-15] or the accuracy [16-18] of responses in neuronal populations processing the relevant sounds. More intricate theories suggest that attention also affects predictive mechanisms in sensory cortices, [19] or that attentional modulation arises as neural networks adapt to specific tasks in various contexts. [20-23]

Recent methodological advances in electrocorticography (ECoG), [15, 24-27] magnetoencephalography (MEG) [11, 28] and electroencephalography (EEG) [25, 29, 30] have revealed that attention enhances neuronal tracking of speech sounds. This amplification is concordant with modulation of both early (i.e., within 100 ms; e.g., [31]) and late (after 100 ms; e.g., [32, 33]) neural response curves to sound envelope changes, consistent with the view that selective attention shifts neuronal processing in low-level auditory and higher-level speech-sensitive regions towards the features of the attended speaker. [24, 31, 34, 35] These methods, however, lack spatial precision. That is, ECoG studies are limited by the extent of the implanted electrodes, while MEG/EEG source localization is relatively inaccurate especially in the case of simultaneously firing neuronal populations. [36] In contrast, functional magnetic resonance imaging (fMRI) provides better spatial resolution, revealing that selective attention to cocktail-party speech modulates information processing in not only low-level auditory regions but also in extensive superior temporal, inferior parietal, and inferior frontal brain regions (e.g., [37-42]). Furthermore, multivariate pattern analyses on cocktail party fMRI data have indicated that neuronal populations that show differential responses during selective attention to speech are distributed globally in disparate cortical regions. [38, 43] Yet, fMRI has limitations in estimating the timing of these modulations. Therefore, some fMRI studies have employed a combination of language modeling and multivariate analysis of fMRI responses to address the temporal limitations of fMRI when tracking continuous speech. [43, 44] However, here we opted for a different approach by utilizing EEG-fMRI fusion. [45, 46] This technique allows us to overcome the spatial limitations of EEG and the temporal constraints of fMRI, enabling us to estimate the spatiotemporal characteristics of selective attention to audiovisual (AV) speech.

In the present paradigm, participants watched video clips of dialogues between two speakers (dialogue stream) with a distracting speech stream played in the background (background stream; Fig. 1a). To increase attentional demands, we modulated the auditory quality of the dialogue stream with noise-vocoding [47] and visual quality in the videos by masking. [48] We also modulated the semantic coherence of the dialogue stream (Fig.1b-c). We employed a fully factorial design where participants performed two different tasks: 1) attend speech task, where the participants attended to the AV dialogue while ignoring the background speech, and 2) ignore speech task, where the participants ignored both the dialogue and the background speech, and instead counted rotations of a white cross presented visually near the mouth of either speaker. This enabled us to study the effect of selective attention (difference between the attend speech task and the ignore speech task) on both the relevant speech stream (dialogue stream) and the irrelevant background (background stream).

**Figure 1.**
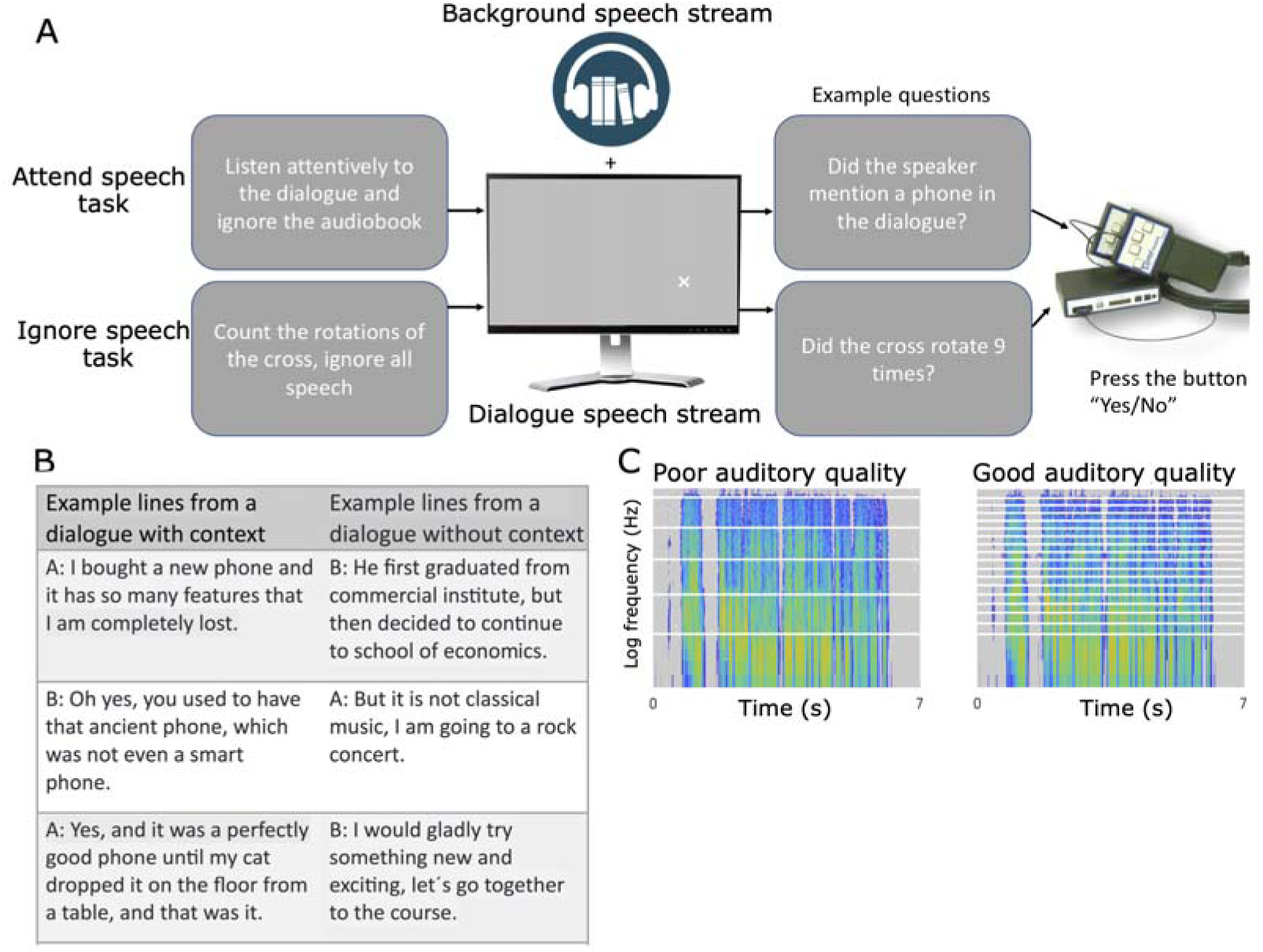
The audiovisual (AV) cocktail party paradigm. **(A)** Participants underwent either EEG (n = 19) or fMRI (n = 19) recordings while watching audiovisual (AV) video clips of dialogues consisting of 7 lines (dialogue stream) with a continuous audiobook (background stream) played in the background. Participants performed two tasks: 1) an attend speech task where they attended to the dialogue while ignoring background speech and 2) ignore speech task where they ignored all speech and counted rotations of a cross presented below the neck of the talker. Dialogues were either semantically coherent or incoherent (B), and the audio quality varied with different levels of noise-vocoding (C). Additionally, visual quality was manipulated with dynamic white noise masking (D).

Using speech envelope reconstruction on the EEG data (Fig. 2), we replicated earlier findings that neuronal tracking is amplified for attended speech (see e.g., [29]). Importantly, however, we found that this amplification was not temporally uniform: the tracking amplitude abated linearly with the procession of the spoken line. Further, neuronal tracking of attended speech displayed nonlinear fluctuations over the course of the dialogue, similar to those previously reported with fMRI. [38] We discuss how such temporal dynamics may arise due to interactions between prediction and attention and other non-linear plastic effects in speech processing circuits. [19, 20] To evaluate the minute temporal modulation of selective attention, we estimated temporal response functions (TRFs) for the EEG data separately for both speech streams (Fig. 3). Finally, we performed EEG-fMRI fusion: Based on representational similarity analysis (RSA) we identified brain regions in the fMRI data that contained representational structures similar to those calculated from TRFs, resulting in a TRF-fMRI correlation time series for each brain region (Fig. 4, Suppl. Video 1 and 2, www.mv.helsinki.fi/home/jkaurama/vdialog/, www.mv.helsinki.fi/home/jkaurama/vbook/). This analysis indicated that attention facilitates recurrent feedforward-feedback loops in the ventral processing stream (see [2]).

**Fig. 2.**
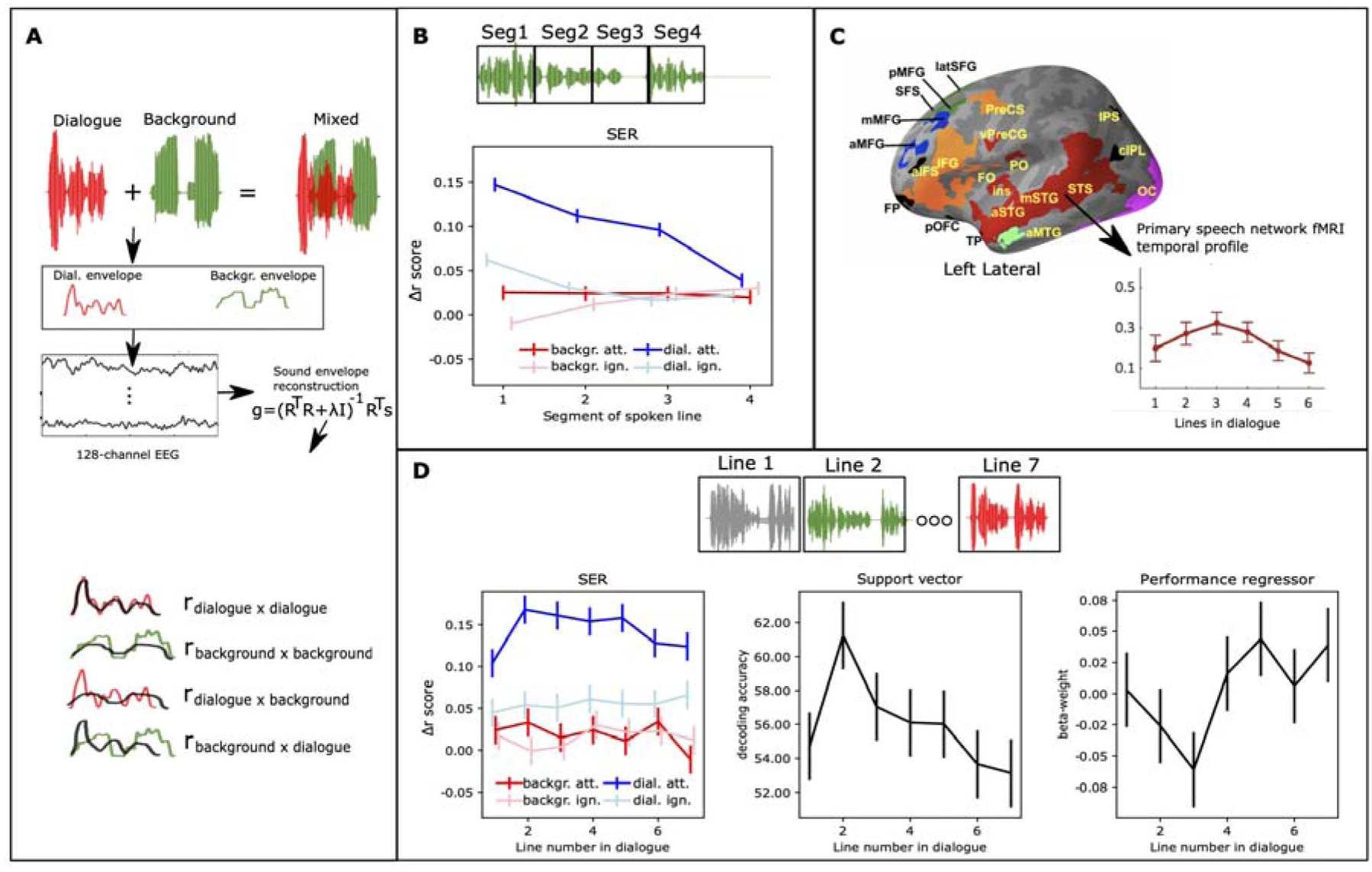
Schematic illustration of speech envelope reconstruction (SER) and SER results. **(A)** Participants heard and saw AV dialogues with overlapping background speech, i.e., a mixed auditory signal. SER was employed to assess neural tracking of the dialogue and background stream. First, we extracted the amplitude envelope for both speech streams. Then using data from all 128 EEG channels, we separately reconstructed the amplitude envelopes for the dialogue and background stream. To assess the accuracy of neural tracking, we correlated the reconstructed speech with its corresponding envelope and compared this to correlations with the opposite envelope. Accuracy values in B-C represent Δ*r* (r-difference scores) between direct correlations and across-reconstruction correlations. **(B)** SER accuracy exhibited a significant linear temporal decrease within each line of the attended dialogue stream. **(C)** Our prior fMRI study [38] demonstrated that attention related modulation changed from line-to-line in a nonlinear fashion. **(D)** SER accuracy displayed a similar non-linear temporal pattern as fMRI (C), but specifically for the attended speech. This trend was observed in both univariate SER accuracy analysis (left) and multivariate support vector machine (SVM) decoding (middle; details in “Decoding analysis of SER accuracies”). Participants’ SER accuracy was predicted based on their behavioral performance for the attended dialogue stream (right), and this prediction (beta-weight) inversely followed SER accuracy. Error bars indicate ± SEMs.

**Fig. 3.**
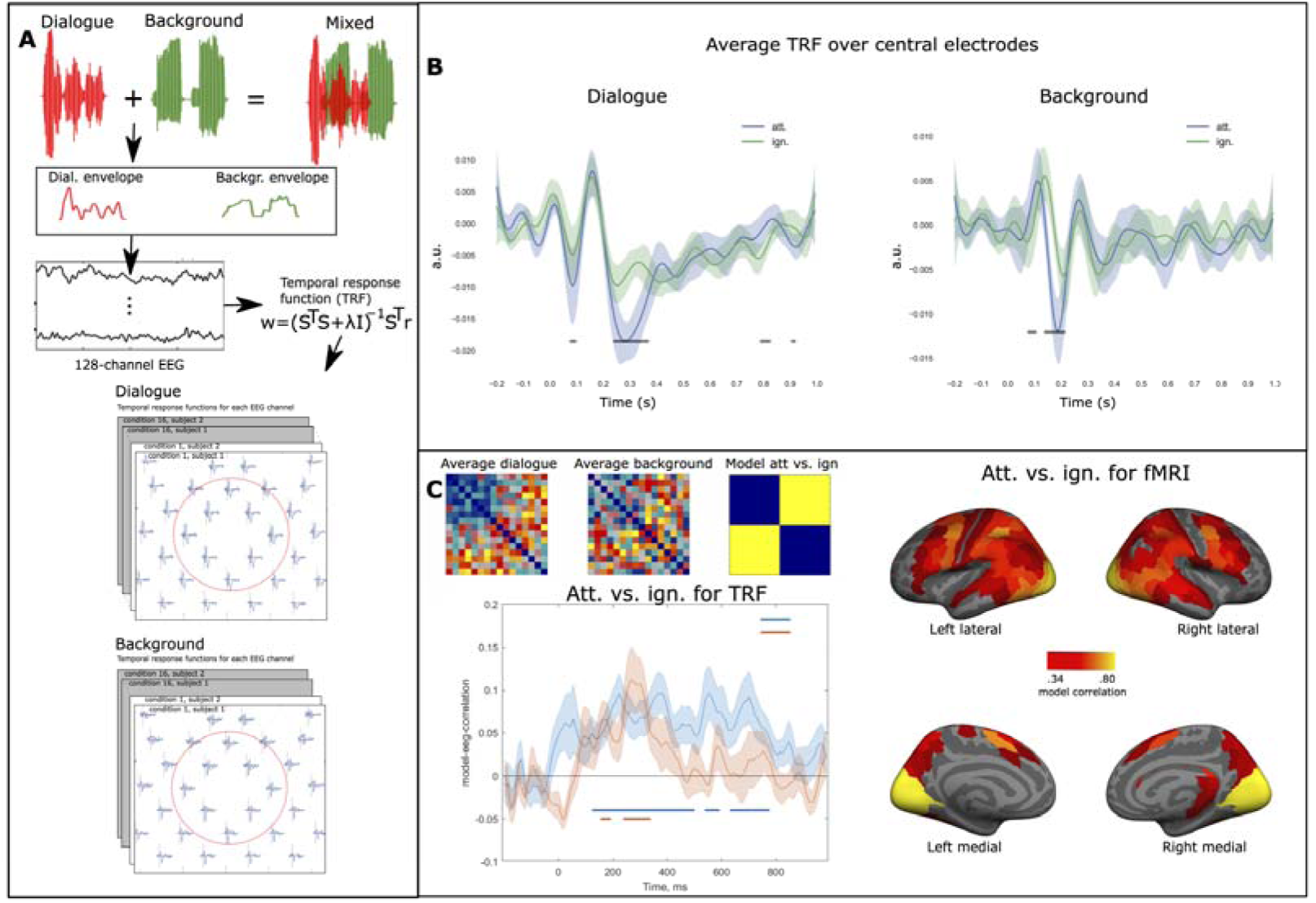
Schematic of temporal response function (TRF) estimation and TRF results. **(A)** TRFs were estimated using the same speech amplitude envelopes as in our SER analysis, separately for the dialogue and background streams. **(B)** Average TRFs over frontocentral electrodes, with points indicating significant differences between the two TRFs (paired permutation *t*-test *df* = 18, note the two streams have separate y-scales). **(C)** Left: Representational dissimilarity matrices (RDMs) were constructed using TRFs for all 16 conditions (first the 8 attend speech conditions and thereafter the 8 ignore speech conditions). This involved pairwise correlations for each condition combination at each time point across EEG channels. The upper left corner shows the average TRF RDMs for both dialogue and background streams. The plot in the left corner displays the correlation between an attentional task model (attend speech vs. ignore speech, att. vs. ign.) and the two TRF RDM timesereies, with significant points displayed below the plot (FDR corrected, one-sample *t*-test, *df* = 19). Right: Similar to TRFs, fMRI RDMs were constructed using searchlight SVM decoding across the 16 conditions, resulting in voxel specific RDMs. Regions with above-average correlations between the attentional task model and fMRI RDMs are displayed (HPC parcellation). Shading indicates ± SEM

**Fig. 4.**
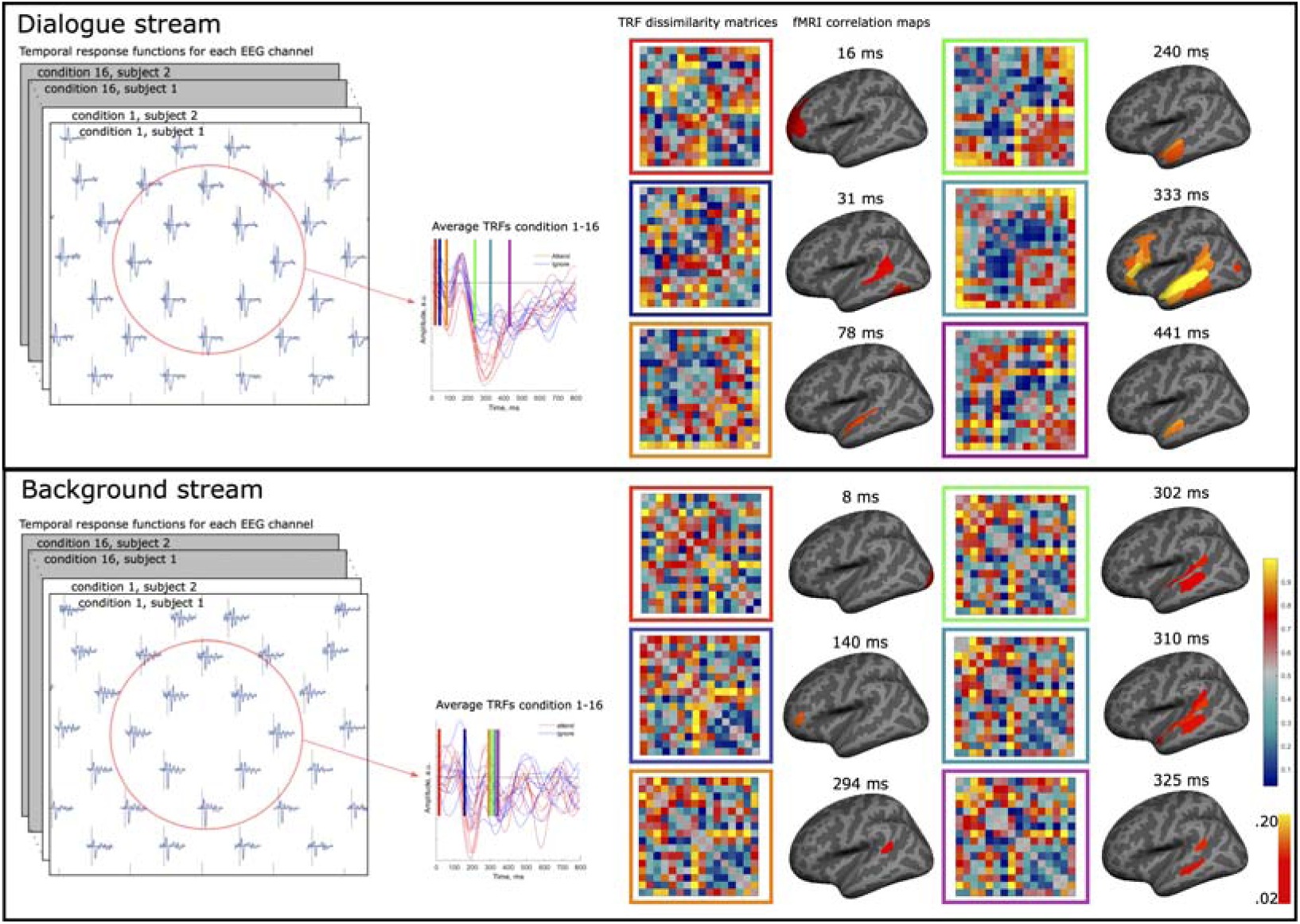
Schematic illustration of TRF-fMRI fusion and results. TRFs were separately estimated for the dialogue (upper part) and background streams (lower part) for each combination of semantic coherence and audiovisual quality and EEG channel. Average TRFs are displayed for frontocentral electrodes in the middle column (attend speech: red, ignore speech: blue). We constructed TRF RDMs for each timepoint by correlating each EEG channel TRF pairwise across conditions and participants. Similar fMRI RDMs were constructed based on SVM decoding between the 16 condition pairs from fMRI data. Thus, we constructed similar RDMs for the EEG and the fMRI, allowing us to fuse information from both datasets by correlating vectorized TRF RDMs with fMRI RDMs, controlling for task and opposite speech stream TRF RDMs (see Fig. 3c). To identify fMRI activations which corresponded to TRF RDMs at different time points, we conducted one-sample t-tests (*df* = 18, FDR corrected) averaged across HCP parcellation ROIs. Six time points of this TRF-fMRI RSA analysis is displayed for both the dialogue (upper part) and background streams (lower part) on the right side of the figure. For the full time series, refer to Suppl. Video 1–2.

## Results

### Attentional modulation of speech envelope reconstruction accuracy fluctuates during the course of the dialogue

We used the accuracy of speech envelope reconstruction (SER) to study how selective attention and our other experimental manipulations affected the neuronal tracking of the two separate speech streams of the presented AV cocktail party speech (i.e., the dialogue stream and the background stream).

In brief, multidimensional transfer functions were estimated based on all EEG channels for the dialogue streams and the background streams separately for each combination of Attentional Task, Semantic Coherence, Auditory Quality, Visual Quality, Line of the speech stream (1–7) and Segment of the line (1–4). Thereafter, the accuracy of the speech reconstruction was assessed by correlating the reconstruction with its corresponding speech envelope and correcting for spurious correlations (see “First-level analysis of EEG-data” and “Univariate analysis of EEG data” for details, and Fig. 2a). The strength of the correlation between the SER and its corresponding speech envelope is generally assumed to reflect the accuracy of neuronal entrainment to the input speech. [24] We used linear mixed models to assess the effects of the repeated factors Attentional task, Semantic Coherence, Auditory Quality and Visual Quality on SER accuracy, separately for the dialogue stream and the background stream.

SER accuracy for the dialogue stream was significantly modulated by attention (*F_1,18.7_ =* 67.2, *p* < .001, η*^2^* = .78). That is, SER accuracy was higher in the task where participants attentively listened to the dialogue streams (mean Δ*r* = .14, SEM = .004) than in the task where they ignored the dialogue stream (mean Δ*r* = .05, SEM = .005). This is in line with previous studies that have shown that selective attention to a specific speech stream strongly increases neuronal tracking of that speech stream compared to the ignored speech streams. [29, 51, 52] Please refer to Suppl. Text and Suppl. Fig. 1–2 for all other significant effects in these linear mixed models and their correspondences to the behavioural performance results.

Next, we analysed whether SER accuracy changed during the dialogue. First, we examined within-line effects, that is, whether SER accuracy changed within the line (lines were divided into four equal length segments). As seen in Fig. 2b, there was a significant linear decrease in SER accuracy throughout the line for the dialogue stream when it was attended, and to some extent for the dialogue stream when it was ignored but not for any other combinations of attentional task and speech stream (Fig. 2b; significant Condition × Segment interaction, *F_9,116_ = 11.2, p* < .001, η*^2^* = .46; linear mixed model with the repeated factors Condition (attend speech task dialogue stream; attend speech task background stream; ignore speech task dialogue stream; ignore speech task background stream) and Segment (1–4)).

In our previous study utilising fMRI, we reported that in the auditory cortex attention related modulations changed in a linear-quadratic fashion during the dialogue (i.e., increased in the beginning of the dialogue and abated thereafter; see Fig. 2c [38]). Neuronal tracking has been suggested to be most strongly affected by neuronal processes in the superior temporal cortex, [29] and thus we expected similar temporal effects here. However, unlike with fMRI, here we could assess whether the previously reported temporal modulations were due to the processing of the attended or the ignored speech stream because they are separable in the EEG data. As seen in Fig. 2d (left), SER accuracy showed a similar temporal profile as previously observed with fMRI. In other words, SER accuracy increased in the beginning of the dialogue and abated towards the end. Further, this temporal effect was only evident when the participants selectively attended to the dialogue stream (Fig. 2d left; significant Condition × Line Number interaction, *F_18,137.3_ =* 2.4, *p* < .002, η*^2^* = .24; linear mixed model with the repeated factors Condition (attend speech task dialogue stream; attend speech task background stream; ignore speech task dialogue stream; ignore speech task background stream) and Line Number (1–7)).

Next, we considered the possibility that the slow temporal effects we found in the SER accuracy data were only evident when analysing SER accuracies separately for each speech stream. That is, it might be that weaker neuronal tracking of the dialogue stream causes a similar concordant change in the neural tracking of the background stream, and thus the contrast between the two streams remained constant throughout the dialogues. Therefore, we performed a multivariate analysis that integrated information from both the dialogue stream and the background stream. Specifically, we assessed whether classification of trials as belonging to the attend speech task or the ignore speech task (using the SER correlations for the dialogue streams and the background streams as input) changed over the course of the dialogue (for details, see “Decoding analysis of SER accuracies”). This analysis revealed that decoding accuracy changed in a similar fashion as SER accuracies of the attended dialogue stream alone (Fig. 2d, middle; linear mixed model with Line Number as the repeated factor and decoding accuracy as the outcome, *F_6,25.6_* = 2.7, *p* < .03, η*^2^* = .39).

Previous studies have shown that SER accuracy correlates positively with behavioural performance. [29] Therefore, attentional lability [53] during different parts of the dialogue could be considered a simple explanation for the slow temporal modulations. This attentional lability should also be observed in the behavioural performance. However, there was no significant change in the behavioural performance in the attend speech task across the lines of the dialogue (generalized linear model with Line Number as a repeated factor; χ*^2^_6_* = 9.8, *p* > .13, see also Suppl. Fig. 2 right, and [38]). Furthermore, unlike previous reports [29], we found no significant general association between SER accuracy in the attend speech task (dialogue stream) and behavioural performance. However, we found that the association between performance and SER accuracy changed over the lines of the dialogue (Fig. 2d right, linear mixed model with Line Number as a repeated factor; *F_6, 28.6_* = 2.9, *p* < .02, η*^2^* = .37). This temporal profile was inverse to the SER accuracy temporal profile. That is, behavioural performance was negatively associated with the lines that showed the highest SER accuracy and positively associated with the lines that showed the lowest accuracy.

### Attention modulates temporal response functions of the attended and ignored speech streams

Speech reconstruction analysis has the advantage of maximizing the power of finding effects in the EEG data, because it integrates information across channels and timepoints to estimate the optimal reconstruction of the sound stimulus. This, however, has the drawback of losing timing and location information in the neural signatures. Therefore, we also performed encoding modelling, separately for each combination of speech stream and listening condition (Fig. 3a). In this model, the speech envelope was used as a regressor in a ridge regressor model, performed separately for data from each EEG channel (see “Univariate analysis of EEG data”). The output of this analysis is a temporal response function (TRF), which describes the convolution in time needed to translate the speech envelope into the EEG data. With some caveats, TRFs can be conceptualised as event-related potentials (ERPs) to a continuous variable. [11] The caveats are that TRFs are filtered more heavily than standard ERPs (we used passband of 0.5–10 Hz) and the choice of regularisation smears exact temporal information, and due to this, the estimated timing of neural events cannot be assumed as exact as for standard ERPs.

We analysed whether selective attention modulates TRFs in frontocentral electrodes (optimal for picking up auditory cortex attention effects; e.g., [54, 55]) separately for the dialogue stream (Fig. 3b, left) and the background stream (Fig. 3b, right). Selective attention significantly enhanced the TRFs for the dialogue stream and this effect was present at two intervals between 0–800 ms, first at ca. 50–100 ms and then at ca. 200–400 ms after sound envelope changes (paired permutation *t-*tests, *df* = 18). This is consistent with previous ERP and TRF studies showing that attention modulates auditory processing of speech relatively early (i.e., within 100 ms), but that the strongest modulation is found at later timepoints. [11, 35, 56-59]

Selective attention also changed both the timing and the amplitude of the TRF to the background stream (at ca. 50–200 ms; paired permutation *t-*tests, *df* = 18). It is important to note that the background stream was ignored in all conditions. However, during the attend speech task, the participants had to actively suppress the background stream, while in the ignore speech task they focused on visual stimuli, designed to automatically keep attention away from all speech streams. Thus, since especially early components of the TRF response likely originate from the auditory cortex [35] it could be expected that the early components of the TRF to the background stream would be smaller for the attend speech task than the ignore speech task. However, our results indicated the reverse. This pattern might arise if participants had involuntary momentary lapses of attention to the wrong speech stream [11] during the attend speech task, causing enhancements also in the background speech stream. We find this unlikely, however, because such lapses would likely cause more variance in the background stream TRFs, rather than the change in amplitude seen in the present results. Furthermore, previous studies using the same paradigm [42] have found that in general, participants do not remember topics of the background stream.

### EEG TRF – fMRI fusion reveals that attention facilitates several feedforward-feedback loops related to the processing of cocktail-party speech

Next, we performed multivariate representational similarity analysis (RSA) on the TRFs. TRFs were estimated for each condition (16 conditions, 8 attend speech task, 8 ignore speech task, for the exact order of the conditions see “Multivariate analysis of TRFs”) and channel (128 channels) separately for the dialogue and the background streams. Thereafter, for each sample of the TRFs (128 Hz, ca. 8 ms samples), we performed pairwise correlations across the EEG channels for each condition pair to construct dissimilarity matrices (*1-r*). This resulted in one TRF representational dissimilarity matrix (RDM) for each timepoint of each speech stream. As can be seen in Fig. 3c (upper left corner), especially in the dialogue stream, the attend speech task conditions are generally similar to each other and dissimilar to the ignore speech task conditions, i.e., there is an effect of attentional task. To test when this effect was significant, we constructed a model matrix for the main effect of attentional task (Fig. 3c (upper left corner)) and correlated this model with the TRF RDMs for each time point (one-sample *t*-test, FDR corrected across time points; for other model correlations see Suppl. Fig. 3). This analysis revealed that selective attention modulated TRFs throughout almost the whole time range from 100 ms (after the sound envelope changes) onwards (Fig. 3c, lower left corner). For the background stream, there were significant correlations with the attentional task model between 150–300 ms.

We also performed the same RSA analysis on our fMRI data that used the same paradigm but different participants. Here dissimilarity matrices were generated based on pairwise searchlight SVM decoding between the 16 conditions (see “Decoding analysis on the fMRI data**”** for details). Fig. 3c (lower, right) shows the regions (averaged for each region of the human connectome project atlas, HCP parcellation; [60]) where the correlation with the attentional task model was above average (i.e., *r > .34*, the threshold for significance was *r* > .08). This analysis shows that information that distinguishes the attend speech task from the ignore speech is contained globally in the brain (see also, [38]) which also probably partly explains why the attentional task model correlated with TRF RDM matrices throughout the time interval.

To gain an understanding of how the TRF RDMs corresponded to the fMRI RDMs, we performed EEG-fMRI fusion, [45, 46] using TRF RDMs estimated using the EEG data and fMRI RDMs (see section “TRF-fMRI fusion**“** for details). This was achieved by correlating each TRF RDM with the fMRI RDMs averaged in each ROI from the HPC parcellation. Because the fMRI RDMs integrate the differences for both the dialogue and background stream, while the TRF RDMs separate these effects, we corrected the TRF-fMRI fusion for the TRF RDMs of the opposite speech stream. That is, the dialogue stream TRF-fMRI fusion was corrected for the background stream TRF RDMs and vice versa. We also corrected for the main effect of attentional task in the TRF-fMRI fusion analysis because this effect was global in both the TRF and the fMRI responses (see above; Fig. 3b-c), and thus masks subtle differences between different regions.

As seen in Fig. 4 (upper right corner) and Suppl. Video 1 (www.mv.helsinki.fi/home/jkaurama/vdialog/) for the dialogue stream, the first significant correlations (one-sample *t*-test, *df =* 18, FDR-corrected) between the TRFs and fMRI RDMs arose at ca. 16 ms after sound envelope changes in dorsolateral and dorsomedial frontal regions. Hereafter, correlations arose at ca. 30 ms in posterior auditory regions and slowly thereafter in anterior auditory cortical regions. After 150 ms correlations arose in the anterior temporal lobe and slowly spread (ca. 250 ms) back to the auditory cortex, frontal and speech processing regions in an anterior– posterior fashion. A second anterior–posterior sweep in the auditory cortex occurred (starting at ca. 450 ms). For the background stream (Fig. 4 bottom right corner, Suppl. Video 2, www.mv.helsinki.fi/home/jkaurama/vbook/), there was initially correlations in the visual cortex. At ca. 140 ms there were correlations in the dorsolateral frontal cortex and then at ca. 300 ms correlations arose in the auditory cortex moving in a posterior–anterior fashion. This effect may relate to suppression of the background stream (cf. [61]). The TRF-fMRI correlation patterns for the background stream were lateralized to the left, while for the dialogue stream it was bilateral, which is consistent with earlier neuroimaging studies. [62]

It is important to note that these TRF-fMRI fusion patterns were not due to a main effect of attentional task (Att. vs. ign.) since this effect was controlled for in the analysis. Furthermore, as can be seen in Suppl. Fig. 3, no other main effect model or interaction model yielded FDR corrected significant results. However, based on the uncorrected results (Suppl. Fig. 3) it seems that the correlations were mostly influenced by interactions between selective attention (Att. vs. ign.) and stimulus features. However, many of the correlation patterns in the TRF-fMRI fusion analysis likely arose due to idiosyncratic differences between the different task conditions at different timepoints of speech processing. For timeseries in different a priori regions of interest please see Suppl. Fig. 4.

## Discussion

Our speech envelope reconstruction (SER) analyses on the EEG data replicated that selective attention enhances neural tracking of attended speech. [25, 29, 30] Similarly, we replicated that attending to a specific speech stream enhances its EEG temporal response functions (TRFs), both at early latencies (ca. 30–150 ms, e.g., [56]) and later latencies (ca. 200–400 ms, e.g., [35]). These findings are consistent with the view that selective attention increases the contrast between attended speech and distracting speech through top-down neural signals, which propagate from higher-level cortical regions to sensory regions and serve to enhance the gain of neurons that process the relevant speech. [12-14] While this is a likely explanation for some of our observations, we find it highly unlikely that this model exhaustively explains how attention modulates sensory processing in the auditory cortex, which we will discuss below.

Although the background stream was always ignored, TRFs for the background stream were both temporally expedited and amplified when participants listened to the dialogue stream compared to when both streams were ignored. Thus, it seems that selective attention not only enhances the processing of relevant speech but also modulates the processing of the actively ignored distracting speech (for similar findings, see [11]). Such modulations might reflect active suppression of auditory cortex neurons processing attributes of distracting speech, which has been suggested as a complementary mechanism to increase the contrast between attended sounds and ignored sounds. [57, 63, 64] Alternatively, the effects may reflect that early processing of the background stream cannot be supressed when attending to speech, [32] attention enhancements partially spread to the background stream [65] or attention fluctuates between the two streams. [11] Later studies utilizing source localisation and/or intracranial measurements could reveal both the spatial and laminar attributes of these effects and the neural populations contributing to them.

RSA analyses of the EEG data for both the dialogue stream and the background stream revealed that selective attention strongly modulated TRFs at several latencies that have not been reported in previous studies. Corroborating this, RSA analyses on the fMRI data showed that attention modulated information processing in extensive cortical fields, not limited to regions associated with speech processing or executive functions. Thus, these results cast doubt on models that highlight simple interactions between frontal and sensory neural networks as origins for selective attentional effects. Rather, our results suggest that selective attention modulates a multitude of different sub-processes widely distributed in the brain (see also [38]).

Using RSA, we performed TRF-fMRI fusion, which showed that attentional modulation of information flow between sensory regions and higher-level regions displayed reliable spatial and temporal characteristics. The earliest modulations were found in the lateral, medial, and inferior frontal cortices at around 8–16 ms. This is consistent with earlier MEG source localisation of attentional effects on speech related TRFs [11] and might reflect preparatory signals biasing the attentional speech processing (e.g., when the quality of the sensory input is poor). Thereafter, information flow generally followed the ventral stream model, [2] with information processing first being modulated in the secondary auditory cortex (around 30 ms), continuing anteriorly to the superior temporal cortex and finally to the anterior temporal lobe (at around 150 ms). At later latencies (after 200 ms) several back-propagating loops of information flow between the anterior temporal cortex, frontal cortex, and the auditory cortex can be discerned. This suggest that information flow during active processing of cocktail-party speech is associated with reverberant bidirectional (feedforward–feedback) informational flow from sensory regions to regions associated with semantic, [8] syntactic [9] and executive functions, [66] within the ventral processing stream.

As previously mentioned, we found that attention enhanced the neural tracking of the attended speech. However, this modulation was not uniform in time, i.e., the SER accuracy linearly decreased within the line of the dialogue (ca. 5 s long). This type of decrease could be explained within a predictive coding framework, [67] assuming that information accumulates as the line proceeds, which constrains prediction error (PE) in neural networks. [51, 68] Importantly, however, we found that decreases in SER were most consistently observed for attended speech. Thus, if the SER temporal profile is explained by predictive mechanisms, such mechanisms seem to depend on selective attention. Indeed, some current models postulate that predictive coding mechanisms and selective auditory attention interact during attentive processing of sensory information (cf. [19, 69, 70]). In the model proposed by Schröger et al.,[19] the attentional processing of relevant sounds is biased in the auditory cortex through recurrent loops, with higher-order processing networks, establishing an ‘attentional trace’ which maximally distinguishes the features of the attended sounds from the features of the irrelevant sounds. In this model, selective attention improves the precision and gain of prediction errors (PEs) generated by neurons encoding the attended stimuli. These enhanced error signals are concurrently sent to regions at the higher level of the processing hierarchy, which in turn send stronger modulatory signals to lower levels of the hierarchy. Thus, attention may influence feedback/feedforward loops, which interact with for example the predictability of the input. This model seems to explain quite well the present linear decrease effects. The model also gives a framework for understanding our TRF-fMRI fusion results, suggesting that the recurrent feedforward/feedback loops reflect the propagation of PE from the lower level of the hierarchy to the next level, on the one hand, and correcting predictive signals from the higher level to the lower level, on the other.

We also found that the strength by which selective attention enhances neural tracking of speech changes on a slow temporal scale (from line-to-line of the dialogue, Fig. 2c). In contrast to the linear decrease seen within a line, the neural tracking first increased up to the middle of the dialogue, and thereafter decreased towards the end of the dialogue. From the predictive coding framework, it could be postulated that such a temporal profile would arise if the ability of attention to maximally increase the gain of PEs takes time to build up, causing an initial increase in SER. The subsequent decrease could be explained, as for the within-line effects, by predictions becoming more stable towards the end of the dialogue. This account, however, fails to explain why there is no indication of such a delay in facilitating attentional processes within the line. Furthermore, based on this account, it would be expected behavioural performance to improve as the dialogue proceeds and the model of the heard speech becomes increasingly accurate. We did not, however, find any evidence for such changes in the behavioural performance data. Importantly, in our previous publication on the fMRI data, we reported similar slow temporal changes of attention related modulations in the superior temporal cortex. [38] In that paper, we suggested that the temporal modulations arose due to recruitment of additional neuronal resources in speech networks that may aid in automatizing speech processing. This account is based on the model proposed by Kilgard, [20] originally used to explain why attention and plasticity initially recruit neurons in the sensory cortex, which after task automatization no longer participate in the task. According to Kilgard, when the task is initially learned, all possible neuronal networks that may be useful to solve the task at hand are recruited. Gradually, the unnecessary, less informative neuronal networks are pruned out, and the most efficient network ends up performing the task (sparse coding). Thus, the slow temporal profiles seen in the current study may reflect that in the sensory cortex, neurons that may help in building the attentional trace are initially recruited and subsequently pruned out to encode information in a maximally sparse manner. This account would also explain the perplexing performance-SER association (Fig. 2c). That is, we found that these behavioural performance predicted SER accuracy negatively in the middle of the dialogue when SER accuracy was strongest and positively when accuracy was weakest. Thus, it may be that behavioural associations were negative in the middle of the dialogue because at this point, neuronal resources processing the speech may not necessarily help in performing the task, while towards the end of the dialogue, unnecessary units are pruned out and the association between SER and performance returns to positive.

Models of the auditory system have generally overlooked how factors like attention and active tasks influence the processing of sounds in neural networks. This oversight relies on the premise that attention simply changes neuronal response gain. Our results, however, highlight that the enhanced neuronal tracking of attended speech is not necessarily uniformly associated with more accurate representation of the attended speech (see e.g., [29]) but changes as a function of time due to predictive and/or other nonlinear plastic mechanisms in sensory cortex. We argue that the approach to selective attention needs to be updated to reflect recent views on how cognition is organised in neural systems (see e.g., [71]). Instead of mechanistic models where higher-level networks enhance gain mechanisms in sensory neurons, attention could be modelled as a collection of temporally changing processes that route activity in distributed neural networks according to behavioural demands. Thus, later multi-and single unit recordings in the auditory cortex could test the hypothesis that attention both changes the gain of neuronal populations and initially recruit neuronal resources that may aid in the performance of the task that are later discarded due to optimisation of task performance.

## Methods

### Data and code availability

The raw data reported in this study cannot be deposited in a public repository due to legal and ethical considerations regarding participant privacy. To request access for anonymized data, contact Dr. Patrik Wikman. In addition, processed datasets derived from these data have been deposited at Open Science Framework under Attention and Memory networks (https://osf.io/agxth/) and are publicly available as of the date of publication. Accession numbers or DOIs are listed in the key resources table.

All original code has been deposited at (https://osf.io/agxth/) and is publicly available as of the date of publication. Access numbers are listed in the key resources table. Further information and requests for resources and reagents should be directed to and will be fulfilled Dr. Patrik Wikman.

### Experimental model and study participant details

#### Participants

EEG data were collected from 20 adult university students at the University of Helsinki and Aalto University (11 females, age range 19–28 years, mean 23.4 years). One participant was excluded due to a technical problem with the EEG data acquisition. fMRI data was collected from a separate sample of adult university students at the University of Helsinki and Aalto University comprising 23 adult participants (14 females, age range 19–30 years, mean 24.3 years). fMRI data were excluded based on pre-established criteria. Two participants were excluded due to excessive head motion (> 5 mm) and two participants due to anatomical anomalies that effected coregistration. Thus, data from 19 participants were used in the analyses. The fMRI data has been previously analysed and published in [38], in the present manuscript the data was analysed differently, yielding previously unreported results, e.g., fusion with the EEG-data. All participants were monolingual native Finnish speakers, and they did not have any self-reported neurological or psychiatric diseases. In addition, they had self-reported normal hearing and normal or corrected-to-normal vision. All participants were right-handed, and this was confirmed by the Edinburgh Handedness Inventory. [72] Each participant provided written informed consent and received a monetary compensation for participation. The experiment was accepted by the Ethics Committee of the Hospital District of Helsinki and Uusimaa.

#### Method details

##### Preparation of stimulus materials

The stimuli comprised dialogues between two (female and male) native Finnish speakers. Written informed consent has been obtained from the individual(s) for the publication of any potentially identifiable images or data included in this manuscript (see also, [38-40, 42]). The dialogue topics were about neutral everyday subjects such as the weather. The dialogues comprised seven lines (ca. 5.4 s of duration) followed by a ca. 3 s break (2.9–4.3 s), resulting in a total length of 55–65 s (mean 59.2 s) for each dialogue. The speakers spoke their lines in an alternating fashion, the female talker started the conversation in half of the video clips.

The original dialogues [42] were recorded so that the talkers sat next to one another with their faces slightly tilted towards each other (see Fig. 1A). For more details on the recordings see [42].

In both the EEG and the fMRI experiment, we used 24 of the original dialogues for the coherent context conditions. The rest of the dialogues were used to construct 24 new dialogues for the incoherent context conditions. These semantically incoherent dialogues were constructed by shuffling lines from different dialogues of the 36 original dialogues. Dialogues were chosen based on the location and posture of the speakers so that there would be minimal visual transition between each line of the shuffled dialogues. Because slight differences in lighting and posture of the speakers, we divided the videos into pools of six videos that were maximally similar. In the semantically incoherent dialogues, each of the five lines were from a separate dialogue, and the remaining two from one dialogue. To secure that all lines were equally unpredictable, we ensured that the two lines from the same original dialogue were separated by at least 4 other lines.

The semantically incoherent dialogues were constructed by first removing the audio stream from the video, whereafter the video image was edited with Adobe Premiere Pro CC software with the morph-cut function (Adobe Inc, San Jose, California, USA). To prevent participants from noticing these changes, the transition from one dialogue to another always occurred on the side where the talker was silent. (see [38] Supplementary video material 1–8; https://osf.io/agxth/). The lighting was edited to fade small differences between the different clips.

Two small grey squares (size 1.5° × 1.5 °) were added to the videos below the faces of the speakers. A white cross (height 0.5 °) was placed in the middle of the square below the face of the talker who was speaking at that given moment. This cross faded out immediately as the talker ended their line and reappeared 1.5 s later. Thus, most of the time there were two crosses present in the video (see [38] Suppl. Video material 1–8; unlike in our experiments, these videos have English subtitles). In the visual control task, the disappearance of the cross indicated that the participant should turn their attention to the other side of the video frame. The cross changed from a plus sign (+) to multiplication sign (×) or vice versa, randomly 9–15 times during each dialogue. The cross rotated only on the side where the talker of the dialogue was speaking. During each of the seven lines, the cross rotated 1–4 times, i.e., every 1.25–2.5 s.

The audio streams were noise-vocoded before adding the audio streams back to the videos. [42] This was achieved by dividing the audio streams into 4 (poor auditory conditions) and 16 (good auditory conditions) logarithmically spaced frequency bands between 0.3 and 5 kHz using Praat software [version 6.0.27, 47] The talkers’ F0 (frequencies 0–0.3 kHz) was unchanged (see [42] for details).

To manipulate the amount of visual speech seen by the participants, we added a dynamic white noise masker onto the speakers’ faces (see [42]).

Finally, the poor and good quality audio files were recombined with the poor and good visual quality videos with a custom Matlab script.

As the final step, we added a continuous background stream to the dialogues. We used a freely available audiobook about cultural history (a Finnish translation of *The Autumn of the Middle Ages* by Johan Huizinga, distributed online by YLE, the Finnish Broadcasting company), read by a female native Finnish professional actor. The F0 of the reader was lowered to 0.16 kHz and the audiobook was low-pass filtered at 5.0 kHz [42].

##### Procedure

The videos, including the dialogue stream and background stream described above, were used in our 16 experimental conditions defined by Attentional Task (attend speech, ignore speech; Att. vs. ign.), Semantic Coherence (coherent, incoherent), Auditory Quality (good, poor) and Visual Quality (good, poor). We presented three runs, each containing eight of the 24 coherent video clips (in all coherent context conditions) and eight of the 24 incoherent video clips (in all incoherent context conditions). Thus, all the participants were presented with all the 48 dialogues. Every other run started with the attend speech task, and every other with the ignore speech task. Within the functional runs, the attend speech task and the ignore speech task were presented in an alternating order. The order of conditions and dialogues presented was pseudorandomized. Because we could not entirely randomise the videos into the 16 conditions per run, we used the Latin square to construct four different versions of the experiment (see Suppl. Table 3 in [38]).

Stimulus presentation was controlled by using Presentation 20.0–22.0 software (Neurobehavioral Systems, Berkeley, California, USA). The auditory stimuli were presented binaurally through insert earphones (Sensimetrics model S14; Sensimetrics, Malden, Massachusetts, USA). Before the experiment, the audio volume was set to a comfortable level individually for each participant. It was approximately 75–86 dB SPL at the ear drum. During EEG the video clips (size 26° × 15°) were presented in the middle of a 24-inch LCD monitor (HP Compaq LA2405x; HP Inc., Palo Alto, California, USA) that was at ca. 40 cm from the eyes of the participant. During fMRI the video clips (size 26° × 15°) were projected onto a mirror attached to the head coil and presented in the middle of the screen. Videos were presented on a uniform grey background. In the middle of each run, there was a break of 40 s. During the break, the participants were asked to rest and focus on a fixation cross (located in the middle of the screen, height 0.5°). The distracting audiobook (presented with a sound intensity 3 dB lower than the voices of the viewed male and female speakers) started randomly 0.5–2 s before video onset and stopped at the offset of the video. The differences in dialogue durations were compensated by inserting periods with a fixation cross between the instruction and the onset of the dialogue, keeping the overall trial durations constant.

##### Tasks

During the attend speech task, the participants were asked to attend to the two speakers having a discussion in the videos while ignoring the background speech. After every dialogue, the participants were presented with seven statements relating to the occurrence of a topic in each line from the dialogue by pressing the ‘Yes’ or ‘No’ button on a response pad with their right index or middle finger. Questions were for example, “Did the boy drop his phone?”, “Was there a cat on the table?”. A new statement was presented every 2 s. After the seven statements, the participants were provided with feedback on their performance (number of correct responses).

During the ignore speech task, the participants were asked to attend to the fixation cross presented in the videos and calculate how many times the cross rotated from a multiplication sign (×) to a plus sign (+) and vice versa. Every time the cross disappeared; the participants were supposed to shift their attention to the other fixation cross on the other side of the frame. The participants were instructed to actively ignore all speech stimuli, i.e., the dialogues and the audiobook. At the end of the video, the participants were presented with seven statements about the rotating cross (“Did the cross turn X times?” the X being between 9 and 15 in an ascending order). As in the attend speech task, the response was given by pressing either the ‘Yes’ or ‘No’ button on a response pad. If the participants were unsure, they were instructed to answer ‘Yes’ to all the alternatives they deemed possible. After the seven statements, the participants received feedback on their performance (number of correct responses).

##### Additional task

After completing the three runs, the participants were presented with an additional run consisting of a single dialogue and one set of seven questions (note only in the EEG-experiment). The dialogue employed in this extra run was the one of the 12 original coherent dialogues that were used to create the 24 incoherent dialogues (i.e., these 12 dialogues had not been seen/heard by the participants in their coherent form in the present experiment). The purpose of this additional run was to evaluate how much the participants processed the semantics of the dialogues they were instructed to ignore during the visual control task. The participants were presented with a dialogue video, and they were told to complete the visual control task and hence ignore the dialogue while counting fixation cross rotations. At the end of the video, they were, however, instructed to answer seven yes-no questions about the lines of the dialogue. The dialogues in this additional run were presented in good auditory and visual qualities and with a coherent semantic context as this was considered the type of conversation that would be the hardest to ignore. This task concluded the experiment. Thus, the additional task was completed by 19 participants participating in the EEG experiment. For results on this task, please see [38].

##### Pre-trial

Before the experiment, all participants practised the tasks. In the practice phase, the participants performed the attend speech task and the ignore speech task, using a coherent dialogue not included in the actual experiment. The dialogue was presented with different auditory and visual qualities.

##### Data acquisition

The EEG data were collected at the Department of Psychology and Logopedics, University of Helsinki, in a soundproof and electrically shielded EEG laboratory. The data were registered separately for each of the three runs of each participant, and the overall duration of the EEG measurements was approximately 1.5 hours per participant. The EEG data were recorded with a BrainVision actiCHamp amplifier (128 channels) and a BrainVision actiCAP snap electrode cap with an actiCAP slim electrode set of 128 active electrodes (Brain Products GmbH, Gilching, Germany). The electrode layout was an extended version of the International 10–20 system, and recording reference was at FCz. The amplifier bandwidth was 0–140 Hz and the sampling rate was 500 Hz. The EEG data were recorded with BrainVision Recorder (version 1.21.0402–1.22.0002; Brain Products GmbH, Gilching, Germany). Electrode impedances were checked prior to recording, and they were below 10 kΩ for most electrodes for most participants. When needed, worsened impedances were enhanced in-between the experimental runs.

For a detailed description of the fMRI acquisition, see. [38] We report the parameters used in brief in Table 1.

**Table 1.**
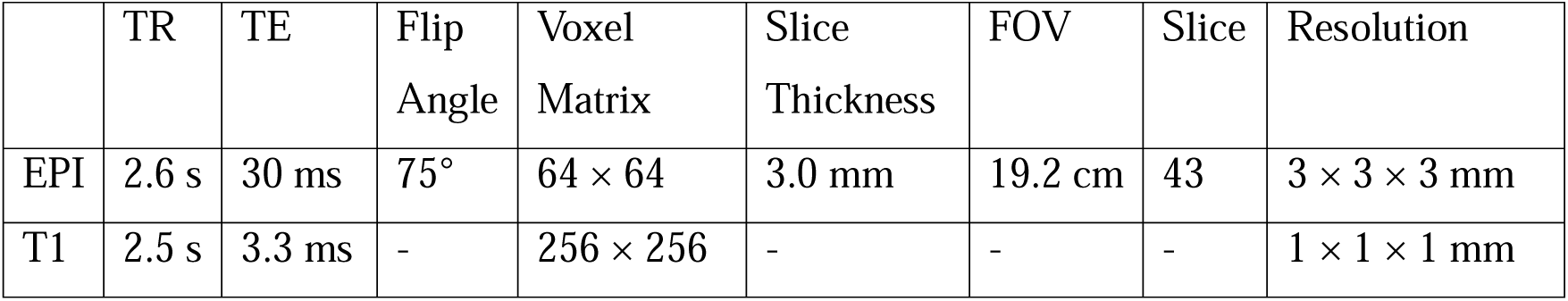
MRI-acquisition parameters used in the fMRI data collection (see [38] for details).

## Quantification and statistical analysis

### Analysis of behavioural data

The total number of questions in the experiment was 336 (48 dialogues × 7 lines). We registered the number of correct answers in each task block. Misses were treated as incorrect button presses. The mean task performance and standard error of mean was used to establish that the participants were performing the task as expected. To analyse participantś performance (EEG/fMRI experiment) during the attend speech and ignore speech task, two separate repeated-measures analyses of variance (ANOVA) were computed with 3 factors: Semantic Coherence (coherent, incoherent), Auditory Quality (good, poor) and Visual Quality (good, poor). ANOVAs were chosen instead of linear mixed models for these analyses to yield comparable results to those reported for the fMRI experiment, which are not reported in the present manuscript, but can be found in [38].

We also analysed the performance line by line to evaluate whether participants’ performance changed during each dialogue (only performed for the attend speech condition performance data gathered in the EEG experiment). Here we used a generalized linear model (identity link function) with the participant added as a random effect (including intercept) and the effect of line was modelled as a categorical repeated measure. The model was run using maximum likelihood estimation with a maximum of 100 iterations to converge.

Statistical analyses were carried out with IBM 18 SPSS Statistics 25 (IBM SPSS, Armonk, New York, USA) software and the results were visualized with Python (Mathworks Inc., Natick, Massachusetts, USA).

### Pre-processing of EEG data

EEG data pre-processing was carried out using the MNE Python 0.22.[73] All channels were referenced to an average reference. Next, the data was manually inspected for channels that would subsequently be interpolated. At least one of the following criteria had to be met for a channel to be chosen for interpolation, and the criterion had to be present persistently throughout at least one of the three experimental runs. The criteria were a flat line response, high-frequency deviation, electrode pop artifacts and body movement artifacts. The deviant channels were temporarily removed from the data.

An independent component analysis (ICA) was fitted on the concatenated runs for each participant separately, using MNE-ICA (picard-type). For each participant, we defined 2–4 components to be denoised that were classified as either blinks, lateral eye movements or heartbeats. Thereafter the raw data from each run was denoised using MNE.ica.apply. After this, the formerly chosen deviant channels were interpolated, the data was bandpass filtered (0.5-10 Hz) and down sampled to 128 Hz for the temporal response function analyses and 64 Hz for the speech reconstruction analyses. Thereafter, the EEG timeseries were cut into 6.5 s epochs based on the dialogue speech trials (see below).

### First-level analysis of EEG data

To estimate the neural response to the two speech streams (dialogue stream and background stream), we performed speech tracking, using both an encoding and a decoding approach. In the encoding approach, we estimated temporal response functions (TRFs) for each EEG channel. In the decoding approach, we reconstructed the speech using data pooled across all 128 EEG channels (speech envelope reconstruction analysis; SER).

The rationale for TRF estimation has been described in detail elsewhere (see e.g.,[74]). In brief, TRFs constitute linear transfer functions describing the relationship between features of the stimulus function (S) and the response function (R; i.e., the EEG channel data). Stimulus features were constructed by extracting sound amplitude envelopes separately for the dialogue stream and the background stream using a Hilbert transform. The envelopes were band-pass filtered (0–10 Hz) and down sampled to 128 Hz for TRFs and 64 Hz for SERs (a lower sampling rate was chosen to speed up analysis for SERs). Thereafter, the envelopes were cut into separate lines (6.5 s) for both sound streams.

In the encoding approach, two separate TRFs were estimated per EEG channel (dialogue and background; Fig. 2a). These TRFs can be conceptualised as a linear composition of partially overlapping neural responses at different time lags (T) to a continuous stimulus, and they are therefore conceptually similar to event-related potentials (ERPs; [11]). We estimated TRFs with the receptive field function (MNE-python: based on the mTRF toolbox which utilises ridge regression), with time lags −200–800 ms, and a common regularisation parameter (A) of 10^5^ ([74]; see Fig. 2a). Note that the regularisation parameter used affects the shape and amplitude of the TRF curves (for simulations, see e.g., [11]) and therefore we chose a common regularisation parameter (based on [74]) and used it in all conditions and participants.

In the decoding analysis, a multidimensional transfer function was estimated using all EEG channels as input (R) in an attempt to reconstruct separately the dialogue stream and the background stream amplitude modulations (see Fig. 3a) using the receptive field function (MNE-python), with time lags (T) of −200–0 ms and a common regularisation parameter (A) of 10^4^.[74] Unlike the encoding analysis, this analysis yields stimulus construction for each time point of the stimulus function (see Fig. 3a).

Both models used a leave-one-out approach, where in each iteration all trials (except one) are selected to train the model (train set), which was then used to predict either the neural response at each EEG channel (TRF) or the speech envelope of speech streams (SER) in the left-out trial (test set). This procedure was repeated with a different train - test partition in each iteration averaged over all iterations.

### Univariate analysis of EEG data

For the TRFs, we tested whether attentional task (i.e., attend speech task vs. ignore speech task) modulated the TRFs averaged across seven frontocentral electrodes (Cz, FCC1h, FCC2h, FC1, FC2, FFC1h, FFC2h; Fig. 3a), separately for the dialogue stream and the background stream for each time bin employing permutation paired *t*-tests (20 000 permutations) using custom scripts written in Python.

For SER, in accordance with [29] we calculated correlations (Pearson) between the original speech stream envelopes and their reconstructions. Thereafter, we cross-reconstruction correlated the stimulus reconstructions and the stimulus envelopes (e.g., correlation between the dialogue speech amplitude and the background speech reconstruction which should be close to zero (Fig. 3a)). Finally, to estimate SER accuracy we used correlation difference scores (Δ*r*; between the correct reconstruction correlations and the cross-reconstruction correlations (Fig. 3b)). For segment level analysis, we divided the stimulus envelopes and the stimulus reconstructions into four segments of equal length and calculated correlations based on these instead of the full line.

The SER accuracies were analysed with different linear mixed models using IBM 18 SPSS Statistics 25. All models included the participant as a random effect and intercept. For repeated factors, the diagonal covariance structure was chosen. If there was more than one repeated factor, a random slope was added for all repeated main effects and interactions using the variance components method. The models were estimated using restricted maximum likelihood estimation with a maximum iteration number of 100 to converge and *df*-estimation was performed using Satterthwaite. Because SPSS does not produce effect size estimates for the fixed effects in linear mixed models, we used the formula partial η^2^ = F × df_1_ / (F × df_1_ + df_2_) [75] to approximate effect sizes were applicable.

To analyse how performance in individual trials affected the dialogue stream reconstruction during the attend speech conditions, we performed a similar two-level analysis commonly used when analysing fMRI data. [38] First, we created separately for each participant a linear regression model with response (correct or incorrect) in each trial as the predictor and SER accuracy for that trial as output. Because there were not enough incorrect responses in any one sub-condition (e.g., coherent, good auditory and good visual quality), all trials were pooled across the eight stimulus conditions. However, because the quality manipulations might affect performance – SER accuracy associations, we added Semantic Coherence, Auditory Quality and Visual Quality as confounds in this model. The β-weight for response was thereafter taken to the second level analysis. The second level analysis was a similar linear mixed model as described above with line entered as the repeated predictor and 5% trimming used for the output variable to remove noise due to the paucity of incorrect trials.

### Decoding analysis on the fMRI data

The pre-processing and first-level analysis pipeline for the fMRI data was the same as that described in detail in [38] (for a brief description see Table 2).

**Table 2.**
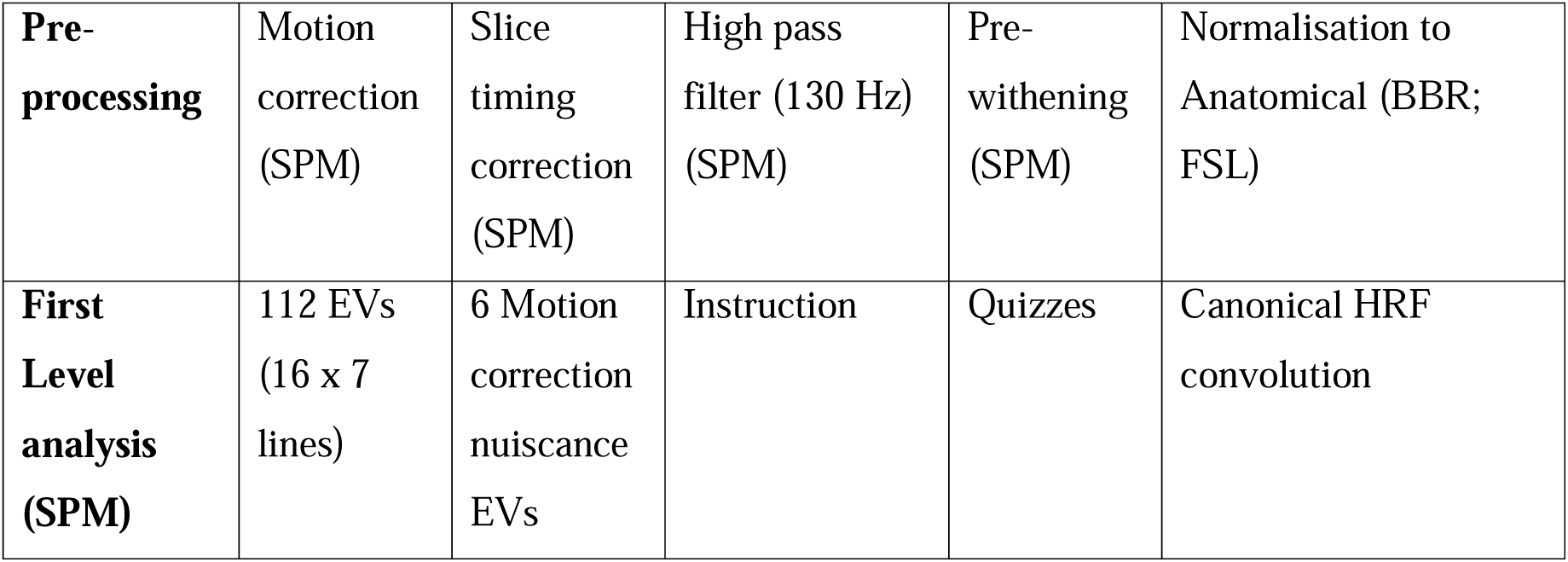
fMRI preprocessing parameters and first-level GLM specifications (see [38] for details).

|  |  |  |  |  |  |
| --- | --- | --- | --- | --- | --- |
| **Pre-processing** | Motion correction (SPM) | Slice timing correction (SPM) | High pass filter (130 Hz) (SPM) | Pre-withening (SPM) | Normalisation to Anatomical (BBR; FSL) |
| **First Level analysis (SPM)** | 112 EVs (16 x 7 lines) | 6 Motion correction nuisance EVs | Instruction | Quizzes | Canonical HRF convolution |

Support vector machine (SVM) decoding with leave-one-run-out cross-validation [76] was used to classify each pair of the 16 conditions (Task (attend speech task, ignore speech task) × Semantic Coherence (coherent, incoherent) × Auditory Quality (good, poor) × Visual Quality (good, poor)) in the fMRI data. Each line constituted an exemplar and each voxel a feature in the analysis. The SVM was conducted with the decoding toolbox [TDT, 77] using the beta images from the first level GLM in the participants’ anatomical space. We used searchlight-based decoding [78] with a radius of 6 mm (isotropic), and with default settings of TDT; L2-norm SVM with regularising parameter C = 1 running in LIBSVM. [79] The resulting accuracy maps for each condition pair were thereafter projected to the Freesurfer average (fsaverage) using the participants’ own Freesurfer surface (surface smoothing: 5mm^2^ full width half maximum smoothing). The pairwise decoding accuracies were averaged within each 360 regions-of-interest (ROIs; HCP parcellation [60]) and representational dissimilarity matrices (RDMs [80]) were constructed for each subject and each ROI. All RDMs in this study are displayed rank-scaled and the conditions are ordered so that the 8 attend speech task conditions are first and the ignore speech task are second. The coherence and quality conditions are in the following order (coherent: co, incoherent: inco, good: g, poor: p, visual quality: v, auditory quality: a; co-gv-ga, inco-gv-ga, co-pv-ga, inco-pv-ga, co-gv-pa, inco-gv-pa, co-pv-pa, inco-pv-pa).

The fMRI RDMs were compared with the attentional task model (Fig. 3c). First, model and data RDMs were vectorised (lower triangular) and then correlated (Spearman *r*) with each other for each ROI and each participant. The statistical significance of the mean correlation above zero was tested with right-tailed *t*-test, FDR-corrected for 360 ROIs.

### Decoding analysis of SER accuracies

Support vector machine (SVM) decoding with leave-one-run-out cross-validation [76] was used to classify SER correlations as either belonging to the attend speech task or the ignore speech task. Each line constituted an exemplar and the four correlations used to define SER accuracies in the univariate analyses (see above) were used as features (*r*: reconstruction of the dialogue stream envelope × the dialogue stream envelope, reconstruction of the dialogue stream envelope × the background stream envelope, reconstruction of the background stream envelope × the background stream envelope, reconstruction of the background stream envelope × the dialogue stream envelope). The SVM was conducted with the decoding toolbox [TDT, 77], using the SER correlations from each participant, with default settings of TDT; L2-norm SVM with regularising parameter C = 1 running in LIBSVM [79], 100 iterations. The resulting accuracies were thereafter analysed using linear mixed models.

### Multivariate analysis of TRFs

RDMs were constructed from the dialogue TRFs as well as background speech TRFs separately for each timepoint by calculating 1-*r* (Spearman) of all conditions across all channels. Like fMRI, the TRF RDMs were compared to model RDMs (See Suppl. Fig. 3). First, model and data RDMs were vectorised (lower triangular) and then correlated (Spearman *r*) with each other for each timepoint and each subject. The statistical significance of the mean correlation above zero was tested with right-tailed t-test, FDR-corrected for 100 timepoints.

### TRF-fMRI fusion

Representational similarity analysis was used to combine EEG and fMRI data. [45, 46] The TRF RDMs for 100 timepoints (0–800 ms) were correlated (Spearman *r*) with the 360 fMRI RDMs. Prior to correlations, the TRF RDMs were averaged across subjects to reduce noise in the data. Furthermore, partial correlation (Spearman *r*) was used, and the effect of task and background speech was controlled for when fusing dialogue TRFs and fMRI, and the effect of task and dialogue was controlled for when fusing background speech TRFs and fMRI. The statistical significance of the mean correlation above zero was tested with right-tailed *t*-tests, FDR-correction was applied for timepoints, ROIs and models (task and dialogue/background speech).

## Author Contributions

PW and KA designed the paradigm, ML wrote the stimulation scripts. Artturi Ylinen, Elisa Sahari, Viivi Kanerva and ES (see acknowledgement) collected the data. PW, ES and VS analysed the data. PW wrote the initial draft of the manuscript. KA and ML applied for funding for the project. All authors edited the manuscript.

## Declarations of interest

The authors declare no competing interests.

## Acknowledgments

This work is supported by the Academy of Finland (grant #297848, “Modulations of brain activity patterns during selective attention to speech”, 2016-2020, KA), and (grant #1348353 “Solving the puzzle of natural auditory object perception - neural mechanisms in humans and animal models”, 2022–2025, PW). We would like to thank Viivi Kanerva, Elisa Sahari and Artturi Ylinen for help with the gathering of the EEG and fMRI data and Ilkka Muukkonen for consultation with the decoding analyses. The funding organisation had no role in the conceptualisation, design, data collection, analysis, decision to publish, or preparation of the manuscript.

Supplementary information is available for this project.

Correspondence and requests for materials should be addressed to Dr. Patrik Wikman.

## Notes

### Competing Interest Statement

The authors have declared no competing interest.

### Summary of Updates

Small changes in text and figures

