## Supplementary document for "Selective attention to audiovisual speech routes activity through recurrent feedback-feedforward loops between different nodes of the speech network"

### Supplementary Text

### Supplementary Speech envelope reconstruction and behavioural performance results.

### Linear mixed models (random intercept) and repeated factors Attention, Semantic Coherence, Auditory Quality and Visual Quality (random slopes for all repeated effects) were performed on SER accuracies separately for the dialogue speech stream and the background speech stream correlations.

### As expected, speech envelope reconstruction (SER) accuracy for the dialogue stream was significantly modulated by attention *(F_1,18.7_* = 67.2, *p* < .001, *η^2^* = .78) because SER-correlations were stronger when participants attended to the dialogue speech streams (mean *Δr* = .14, SEM = .004) than when they ignored the dialogue speech stream (*Δr* = .05, SEM = .005). There were no significant effects of attention on the SER-correlations for the background speech stream.

### There were significant two-way interactions: between Attention and Semantic Coherence (F_1,23.5_ = 7.4, *p <* .01, *η^2^* = .24 ) because, when the participants attended to the dialogue speech stream, SER accuracies were stronger for semantically coherent dialogues than incoherent dialogues; and between attention and auditory quality (*F_1,21.6_* = 5.4, *p <* .03, *η^2^* = .20) because, when participants attended to the dialogue speech, SER accuracies were stronger for dialogues with good auditory quality than dialogues with poor auditory quality. There was also a significant four-way interaction between all factors (*F_1,24,.6_ =* 5.8, *p <* .02, *η^2^* = .19; see Suppl. Fig. 1).

### For the background speech stream there was a significant main effect of Auditory Quality (*F_1,24.6_ =* 5.6, *p <* .03, *η^2^* = .19), because SER accuracies were stronger when the dialogue stream was presented with good (*Δr* = .03, SEM = .004) than poor auditory quality ( *Δr* = .009, SEM = .004); and a significant main-effect of Visual Quality (*F_1,26.9_ =* 7.6, *p <* .01, *η^2^* = .22), because SER-correlations were stronger when the dialogue speech stream was presented with poor quality( *Δr* = .03, SEM = .004) than good quality (*Δr* = .007, SEM = .004).

### Analysis of the performance in the EEG-experiment for the attend speech task yielded partially similar results as the SER analyses. There were significant main effects of Semantic Coherence (*F_1,18_* = 109.0, *p <* .001, *η^2^* = .85),), Auditory Quality (*F_1,18_ =* 30.6, *p <* .001, *η^2^* = .63) and Visual quality (*F_1,18_ =* 28.6, *p <* .001, *η^2^* = .61). All effects were due to performance being better with coherent and good quality dialogues (see Suppl. Fig. 2, left). There was also a significant three-way interaction between the factors (*F_1,18_* = 4.6, *p <* .04, *η^2^* = .2). However, a comparison of Suppl. Fig 1 and Suppl. Fig. 2 reveals that this interaction arose due to different effects in the behavioural performance compared to the SER-correlations. There were no significant differences in performance between the coherence and quality conditions during the ignore speech task (Suppl. Fig. 2 middle).

### Supplementary figures

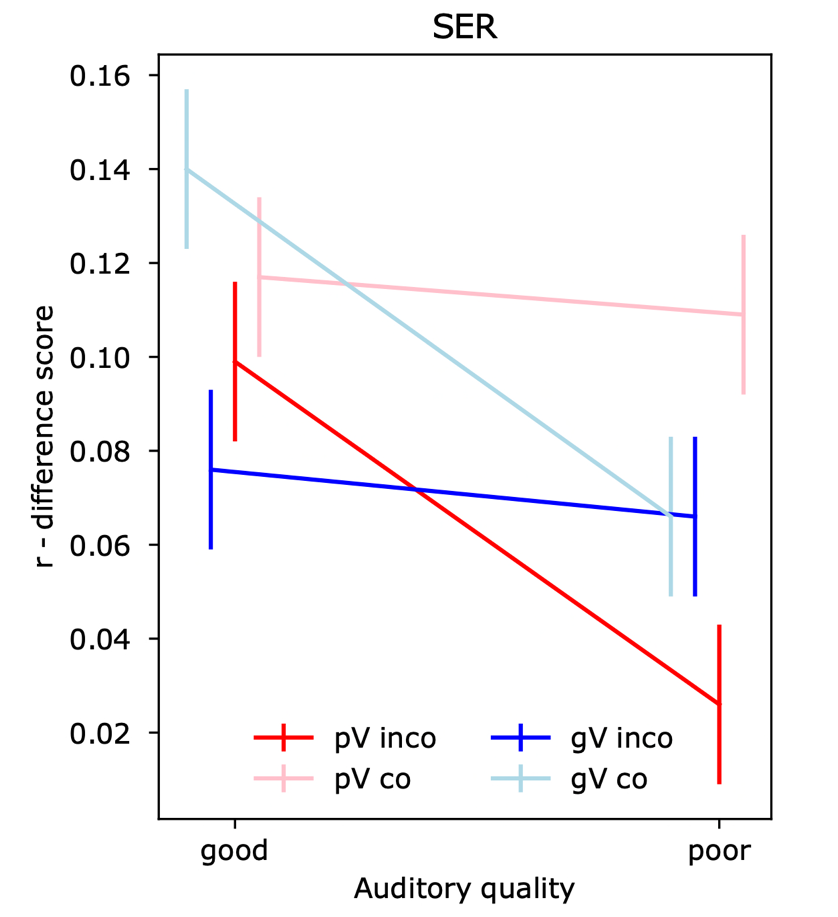

### Supplementary Figure 1. SER-accuracy changed depending on Attentional task, Semantic Coherence, Auditory and Visual Quality. SER-accuracy was estimated using SER-correlation difference scores (see 4.9.4, for details; here the difference between the attend task and the ignore task is displayed). Error bars denote ± SEM. Abbrievations: p, poor; g, good; inco, incoherent; co, coherent; V, visual quality.

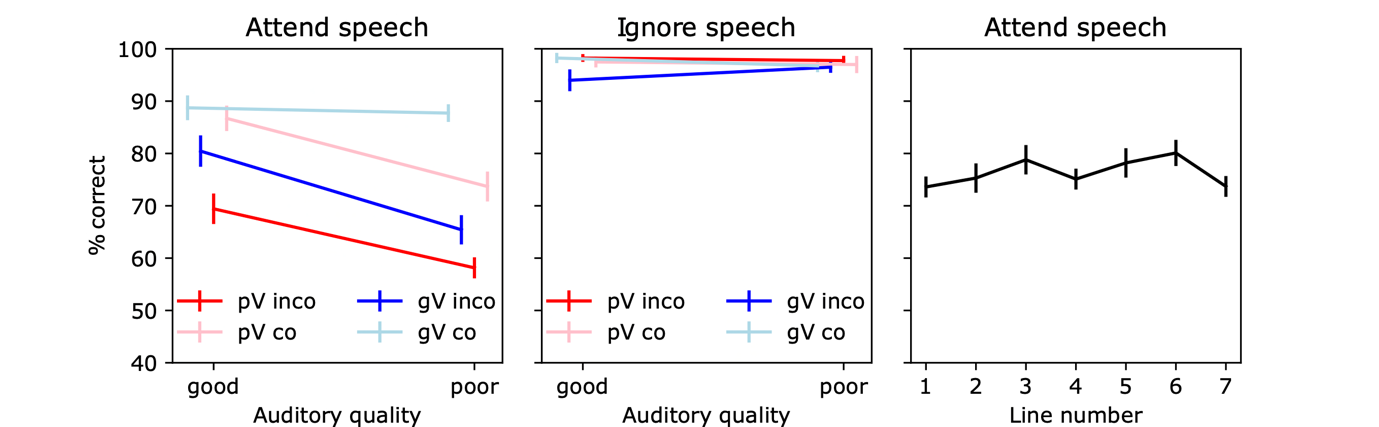

### Supplementary Figure 2. Behavioural results. Performance (percentage of correct answers) for the attend speech task (left), ignore speech task (middle). The rightmost plot shows the percentage of correct answers for the attend speech task by number of lines in the dialogue. Error bars denote ± SEM. Abbrievations: p, poor; g, good; inco, incoherent; co, coherent; V, visual Quality.

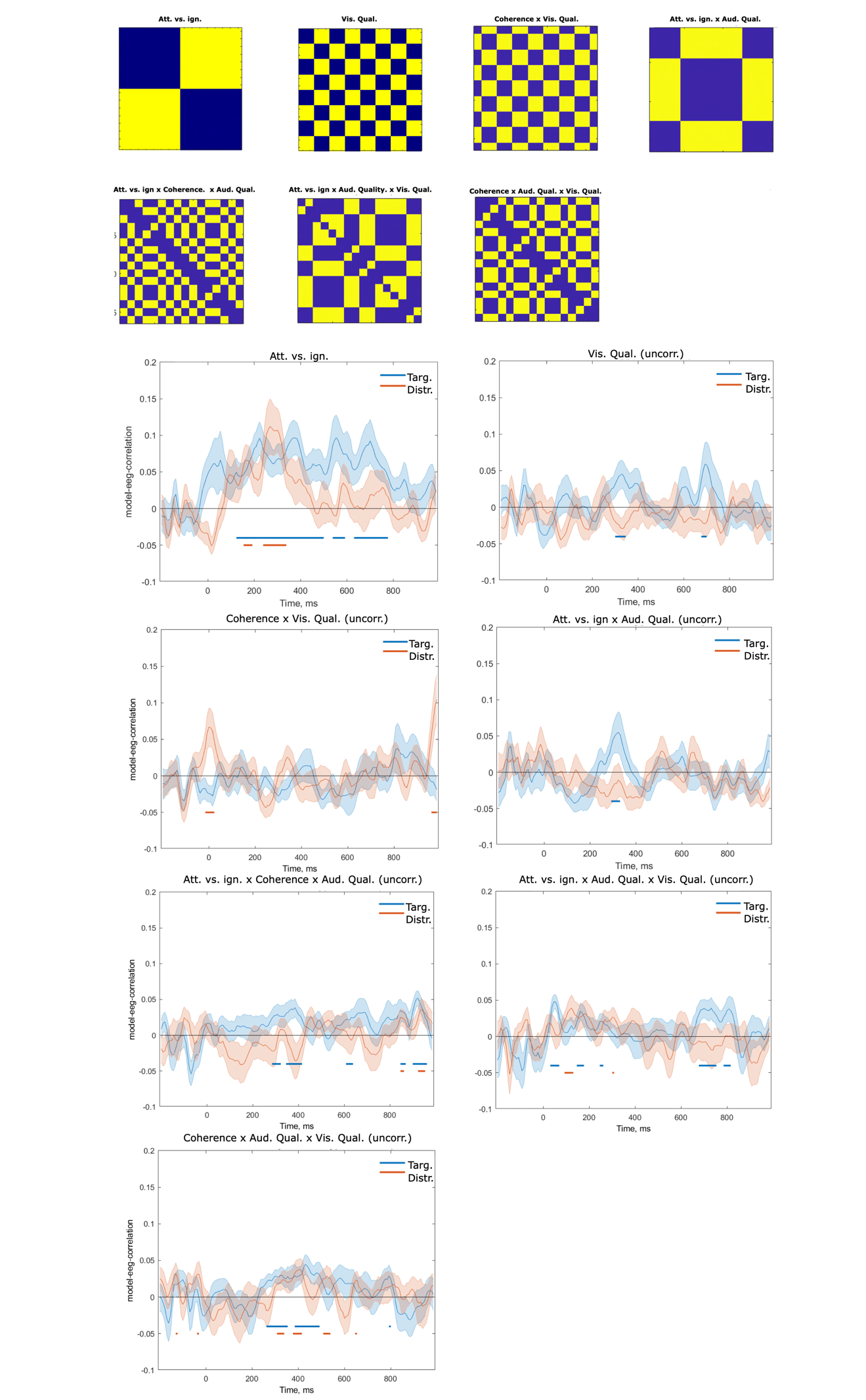

**Supplementary Fig. 3. Representational dissimilarity (RDM) model correlations for the temporal response function (TRF) RDM timeseries separately for dialogue and background speech streams.** We constructed all possible main-effect and interaction RDM models and correlated them with the speech stream RDM-timeseries (see section 4.11). The attentional task (Att vs. ign) model yielded significant correlations (FDR corrected) for both the dialogue and the background stream RDM timeseries. No other model (including those not displayed here) yielded significant FDR corrected correlations (here we display some of the models that yielded reliable uncorrected, p < .05 correlation).

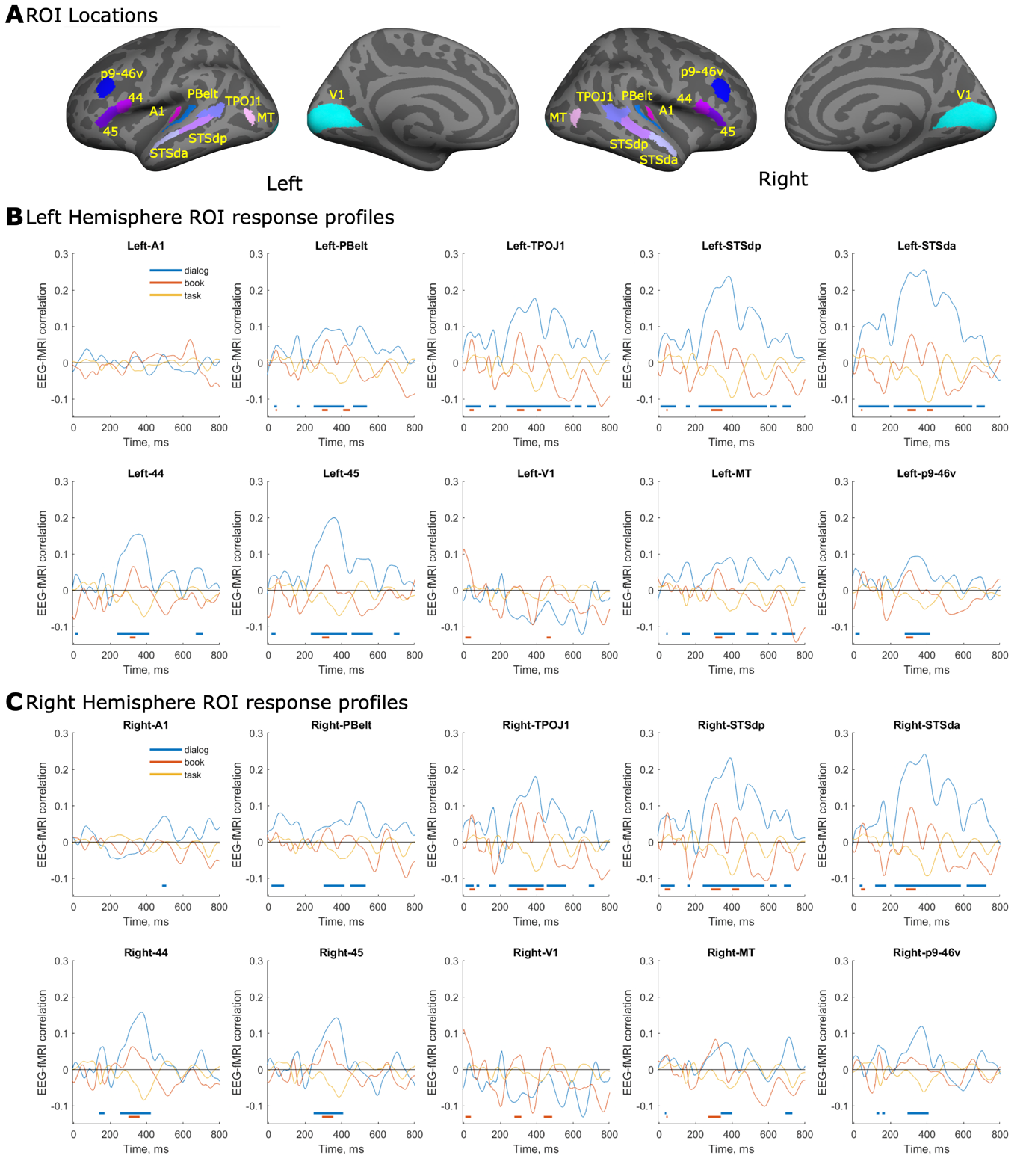

**Supplementary Figure 4. TRF-fMRI fusion results averaged for 10 different a priori selected regions of interest (ROIs).** We performed TRF-fMRI fusion to reveal when there were correspondences between the information structure of the TRFs and the fMRI data (for details see 4.7.8 and 2.3 of the main-text, Fig. 4 and Suppl. Video 1-2). Here we display the TRF-RDM-fMRI correlation timeseries averaged in 10 a priori selected ROIs of the HCP parcellation (For ROI names see ^2^). We selected A1 and PBelt based on the meta-analysis on selective attention effects for speech stimuli in the auditory cortex. ^3^ The STS ROIs (TPoJ1, STSdp, STSda) were chosen because these regions showed the strongest effects of selective attention in. ^1^ The IFG ROIs (44, 45) were selected because they were described as central nodes of the speech control networks in our previous fMRI studies. ^1,4-6^ V1 and MT was chosen because the presented speech was audiovisually presented, i.e., contained moving visual speech. The dorsolateral prefrontal region p9-46v was chosen because this region was shown to change its connectivity with the auditory cortex during selective attention to speech in our previous analyses on the fMRI data.^1^

**Supplementary tables**

|  | TR | TE | Flip Angle | Voxel  Matrix | Slice Thickness | FOV | Slice | Resolution |
| --- | --- | --- | --- | --- | --- | --- | --- | --- |
| EPI | 2.6 s | 30 ms | 75° | 64 × 64 | 3.0 mm | 19.2 cm | 43 | 3 × 3 × 3 mm |
| T1 | 2.5 s | 3.3 ms | - | 256 × 256 | - | - | - | 1 × 1 × 1 mm |

### Supplementary table 1. MRI-acquisition parameters used in the fMRI data collection (see ^1^ for details).

| **Pre-**  **processing** | Motion correction  (SPM) | Slice timing correction  (SPM) | High pass  filter (130 Hz)  (SPM) | Pre-withening (SPM) | Normalisation to  Anatomical (BBR; FSL) |
| --- | --- | --- | --- | --- | --- |
| **First**  **Level**  **analysis**  **(SPM)** | 112 EVs  (16 × 7 lines) | 6 Motion  correction  nuiscance  EVs | Instruction | Quizzes | Canonical HRF convolution |

### Supplementary table 2. fMRI preprocessing parameters and first-level GLM specifications (see ^1^ for details).

1.	Wikman, P., Sahari, E., Salmela, V., Leminen, A., Leminen, M., Laine, M., and Alho, K. (2020). Breaking down the cocktail party: Attentional modulation of cerebral audiovisual speech processing. NeuroImage, 117365.

2.	Glasser, M.F., Coalson, T.S., Robinson, E.C., Hacker, C.D., Harwell, J., Yacoub, E., Ugurbil, K., Andersson, J., Beckmann, C.F., and Jenkinson, M. (2016). A multi-modal parcellation of human cerebral cortex. Nature *536*, 171-178.

3.	Alho, K., Rinne, T., Herron, T.J., and Woods, D.L. (2014). Stimulus-dependent activations and attention-related modulations in the auditory cortex: A meta-analysis of fMRI studies. Hearing Research *307*, 29-41. 10.1016/j.heares.2013.08.001.

4.	Leminen, A., Verwoert, M., Moisala, M., Salmela, V., Wikman, P., and Alho, K. (2020). Modulation of brain activity by selective attention to audiovisual dialogues. Frontiers in Neuroscience *14*, 436. <https://doi.org/10.3389/fnins.2020.00436>.

5.	Wikman, P., Ylinen, A., Leminen, M., and Alho, K. (2022). Brain activity during shadowing of audiovisual cocktail party speech, contributions of auditory–motor integration and selective attention. Scientific Reports *12*, 18789.

6.	Ylinen, A., Wikman, P., Leminen, M., and Alho, K. (2022). Task-dependent cortical activations during selective attention to audiovisual speech. Brain Research *1775*, 147739.
